# Genetic predisposition towards multicellularity in *Chlamydomonas reinhardtii*

**DOI:** 10.1101/2024.08.09.607418

**Authors:** I-Chen Kimberly Chen, Shania Khatri, Matthew D. Herron, Frank Rosenzweig

## Abstract

The evolution from unicellular to multicellular organisms facilitates further phenotypic innovations, notably cellular differentiation. Multiple research groups have shown that in the laboratory simple, obligate multicellularity can evolve from a unicellular ancestor under appropriate selection. However, little is known about the extent to which deterministic factors like ancestral genotype and environmental context influence the likelihood of this evolutionary transition. To test whether certain genotypes are predisposed to evolve multicellularity in different environments, we carried out a set of 24 evolution experiments each founded by a population consisting of 10 different strains of the unicellular green alga *Chlamydomonas reinhardtii*, all in equal proportions. Twelve of the initially identical replicate populations were subjected to predation by the protist *Parmecium tetraurelia* while the other 12 were subjected to settling selection by slow centrifugation. Population subsamples were transferred to fresh media on a weekly basis for a total of 40 transfers (∼600 generations). Heritable multicellular structures arose in four of 12 predation-selected populations (6 multicellular isolates in total), but never in the settling selection populations. By comparing whole genome sequences of the founder and evolved strains, we discovered that every multicellular isolate arose from one of two founders. Cell cluster size varied not only among evolved strains derived from different ancestors but among strains derived from the same ancestor. These findings show that both deterministic and stochastic factors influence whether initially unicellular populations can evolve simple multicellular structures.

## Introduction

A longstanding question in population biology centers around the relative roles that stochastic processes (e.g., mutation, recombination and genetic drift) versus deterministic process (e.g., natural selection) play in generating evolutionary trajectories and outcomes. In many instances of convergent evolution, the deterministic process of natural selection seems to prevail (1–3). For example, on each of the Caribbean’s four largest islands (Puerto Rico, Jamaica, Cuba and Hispaniola), lizards of the genus *Anolis* exhibit four basic morphologies: slender grass-dwelling forms with long tails, ground-dwelling forms with long legs, short-legged forms that creep on twigs, and canopy-dwelling forms with large toe pads. These four body types independently adapted to similar habitats on each island, making it one of the best-studied examples of how evolutionary outcomes can be predictable (2). On the other hand, stochastic processes, such as a lineage’s ancestral history and the levels of variation maintained by mutation and drift, can make evolutionary outcomes largely unpredictable (4). For instance, in *Escherichia coli* and *Salmonella typhimurium* resistance to the antibiotic streptomycin caused lower rate of protein synthesis, thereby reducing the fitness of cells when the drug was not present (5, 6). This fitness cost can be diminished by compensatory mutations at other sites in the bacteria’s genomes (7, 8). However, when many of those compensatory mutations were introduced into wild-type, streptomycin-sensitive backgrounds, they were found to be detrimental, indicating that the fitness effect of those compensatory mutations depended on genetic backgrounds (5, 6, 9). Had those compensatory mutations occurred without the prior streptomycin-resistant mutations, their evolutionary trajectories and fitness outcomes could have been very different.

Laboratory microbial evolution has proved to be a useful tool for investigating under what conditions stochastic versus deterministic factors exert the stronger influence on adaptation evolution. Well-controlled, highly replicated experiments can be carried out for hundreds or even thousands of generations under different types and/or intensities of selection. Such studies have repeatedly shown that similar environments tend to select for similar phenotypes, suggesting that evolution can be deterministic and predictable. For example, all 12 lines in Richard Lenski’s long-term evolution experiment (LTEE) in *E. coli* evolved faster growth and larger cell size than the ancestor (10), as well as similar overall patterns of gene expression (11). In asexual haploid *Saccharomyces cerevisiae* populations, sterility and whole genome duplication repeatedly evolves (12, 13). Of course, stochastic processes like mutations can produce these predictable results, provided that the genetic architecture of a selectively advantageous phenotype is sufficiently complex that it can be approached via different mutational trajectories (14). But sometimes, stochastic events like mutations can produce wholly unexpected outcomes. For example, in one of the twelve LTEE *E. coli* populations that had evolved for >31,000 generations in glucose-limiting, citrate-buffered medium, a weakly beneficial mutation followed by a refining mutation together conferred the ability to exploit citrate (Cit^+^) as a growth substrate. These mutations were strongly selected, resulting in a Cit^+^ dominated, Cit^+^/Cit^-^ population (15–17). A diagnostic feature of wild-type *E. coli* is its inability to ferment citrate under oxic conditions. Thus, this two mutation combo, a tandem duplication that captured an aerobically expressed promoter for the expression of a previously silent citrate:succinate antiporter gene, *citT*, and a promoter mutation that activated expression of C_4_-dicarboxylate:H^+^ symporter *dctA*, brought about a key evolutionary innovation (17). This innovation, which was contingent on an earlier mutation in the citrate synthase gene *gltA*, benefited from even earlier mutations that increased acetate production after the optimization of glucose metabolism, fundamentally altered the ecological and evolutionary trajectory of the affected population (15, 18).

While the roles that stochastic versus deterministic factors play in producing microevolutionary outcomes have been assessed in the lab, it is less clear how such factors have impacted macroevolutionary outcomes, including major evolutionary transitions in the history of life (19). Among these transitions is the evolution of multicellular life-forms from unicellular ancestors, a transition that has occurred repeatedly across the Tree of Life (20–23), and that has served as a gateway to further innovations, such as division of labor among cells, including germ-soma differentiation (19). Historically, our understanding of the mechanisms that have driven this transition has derived chiefly from paleontology (24–26), comparative morphology (27–29), and genomics (30–37), all of which are retrospective in nature. However, in recent years a number of investigators have adopted a prospective approach, and shown that simple, heritable, obligate multicellularity can evolve under selection in the laboratory (38–42). Here, using laboratory evolution we investigate the extent to which two different deterministic processes, selection mode and genetic ancestry, enable a unicellular eukaryote, the green alga *Chlamydomonas reinhardtii*, to make the evolutionary transition from single cells to multicellularity.

*C. reinhardtii* is a member of the volvocine algae (Division Chlorophyta), a group that encompasses species exhibiting different levels of complexity that range from single-celled *C. reinhardtii*, to colonial *Gonium pectorale*, to fully differentiated *Volvox carteri*. Two prior evolution experiments generated heritable multicellularity in *C. reinhardtii* via selective regimens that favor increased size. Using settling selection, Ratcliff, Herron (41) evolved amorphous multicellular clusters of >10^2^ cells bound to one another by a transparent extracellular matrix. Using predation selection by the protist *Paramecium tetraurelia*, Herron, Borin (42) evolved a variety of stereotypical multicellular structures, consisting of 2^2^ - 2^5^ cells surrounded by a common cell wall. These structures bear a striking resemblance to morphologies exhibited by some of the more ‘primitive’ multicellular members of the volvocine algae, e.g., *Pandorina* and *Yamagishiella*.

Unfortunately, instead of starting from the same ancestral strain, the ancestors used in these two selection experiments consisted of different outbred populations, making it impossible to establish whether it was ancestral genotype or the selective regimen that determined which form of multicellularity evolved. To test whether certain genotypes are predisposed to evolve multicellularity in certain forms, or if particular environments select for specific multicellular phenotypes, we carried out two sets of evolution experiments each founded by a population consisting of 10 different strains of *C. reinhardtii*, all in equal proportions. Twelve initially identical replicate populations were subjected to predation by *P. tetraurelia* and 12 replicate populations were subjected to settling selection. Subsamples of these populations were transferred to fresh media every week for a total of 40 transfers (∼600 generations). Because the medium used in this experiment contains abundant nitrate, we expect all reproduction to be asexual (the entry into the sexual cycle is controlled by nitrogen deprivation in *C. reinhardtii*, (43)). Founder strains included two lab strains and eight field isolates, each of which was phenotypically (e.g., cell sizes and chlorophyll content) and genetically different from the others (44). In particular, each of our founder strains contained unique stable polymorphisms that serve as markers for identifying the probable ancestor(s) of evolved multicellular strains.

In our experiments, after ∼600 generations, heritable, obligate multicellular structures arose in four of 12 predation-selected populations, but never in the settling rate selected populations. We found that only two of the original ten strains were ‘parents’ for these multicellular descendants, suggesting that certain genetic backgrounds may predispose those strains to evolve multicellularity under this selective regimen. However, the size and structure of the multicellular types were different even in isolates originating from the same ‘parent’ strain, indicating that stochastic processes like mutation accumulation can lead to strikingly different multicellular forms in populations evolved under the same condition. Together these findings show that both deterministic and stochastic factors can play important roles in shaping the path by which populations evolve simple multicellularity.

## Materials and Methods

### Experimental evolution

A list of the *C. reinhardtii* founder strains used in this study is provided in **Table 1**. Strains were selected on the basis of their geographic isolation from one another, their phenotypic and phylogenetic diversity, and the fact that the genome of each strain has been sequenced (44). In this study, each founder strain was grown to stationary phase (∼6-8 x 10^6^ cells/mL) in TAP liquid medium (45); equal volumes of those cultures were mixed and used as the master stock to start the selection experiment. Twelve replicate populations were subjected to predation selection by *Paramecium tetraurelia* (Carolina Biological Catalogue #131560) as described by Herron, Borin (42) and 12 replicate populations were subjected to settling rate selection (41) (**Figure 1**). Each population was cultured on a 14:10-hour light:dark cycle at 22.5 °C in 15 mL of COMBO medium. COMBO is a fresh water medium suitable for co-culturing algae and zooplankton (46). We used COMBO for both predation and settling rate selection experiments so that the two treatments could be compared. Subsamples of each population were transferred to fresh COMBO medium every week for a total of 40 transfers, with approximately 15 generations elapsing between transfers. For experiments under predation selection, 1.5 mL of co-cultured *C. reinhardtii* and *P. tetraurelia* were transferred to fresh medium every week. For the settling rate selection, each population was transferred to a 15 mL Falcon tube and centrifuged at 100 x *g* for 10 s and only the bottom 0.25 mL was transferred to fresh medium. Under both types of selection, the bottleneck size at each transfer was approximately 1-2 x 10^6^ cells.

**Table 1.**
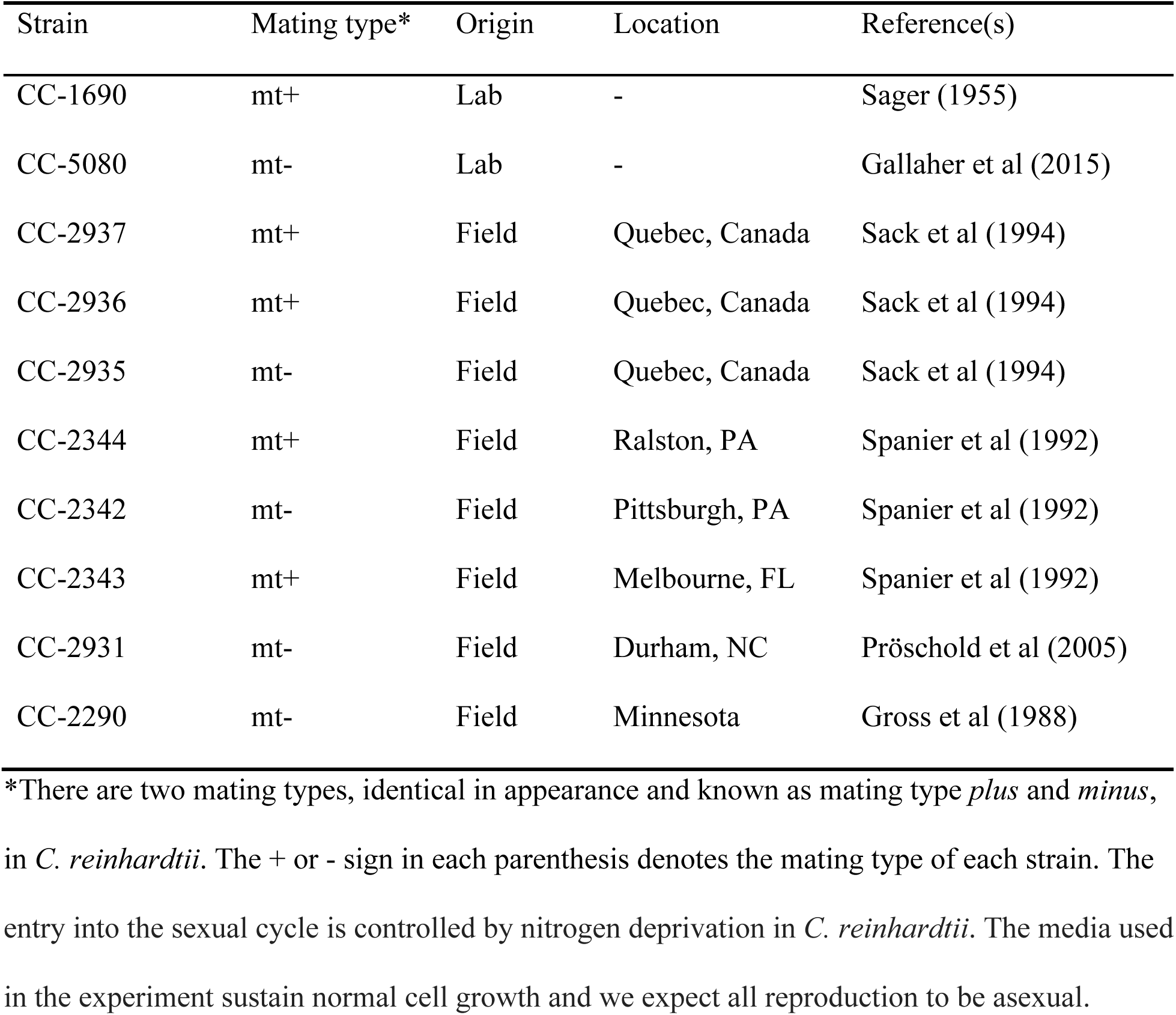
*C. reinhardtii* founder strains used in this experiment.

**Figure 1.**
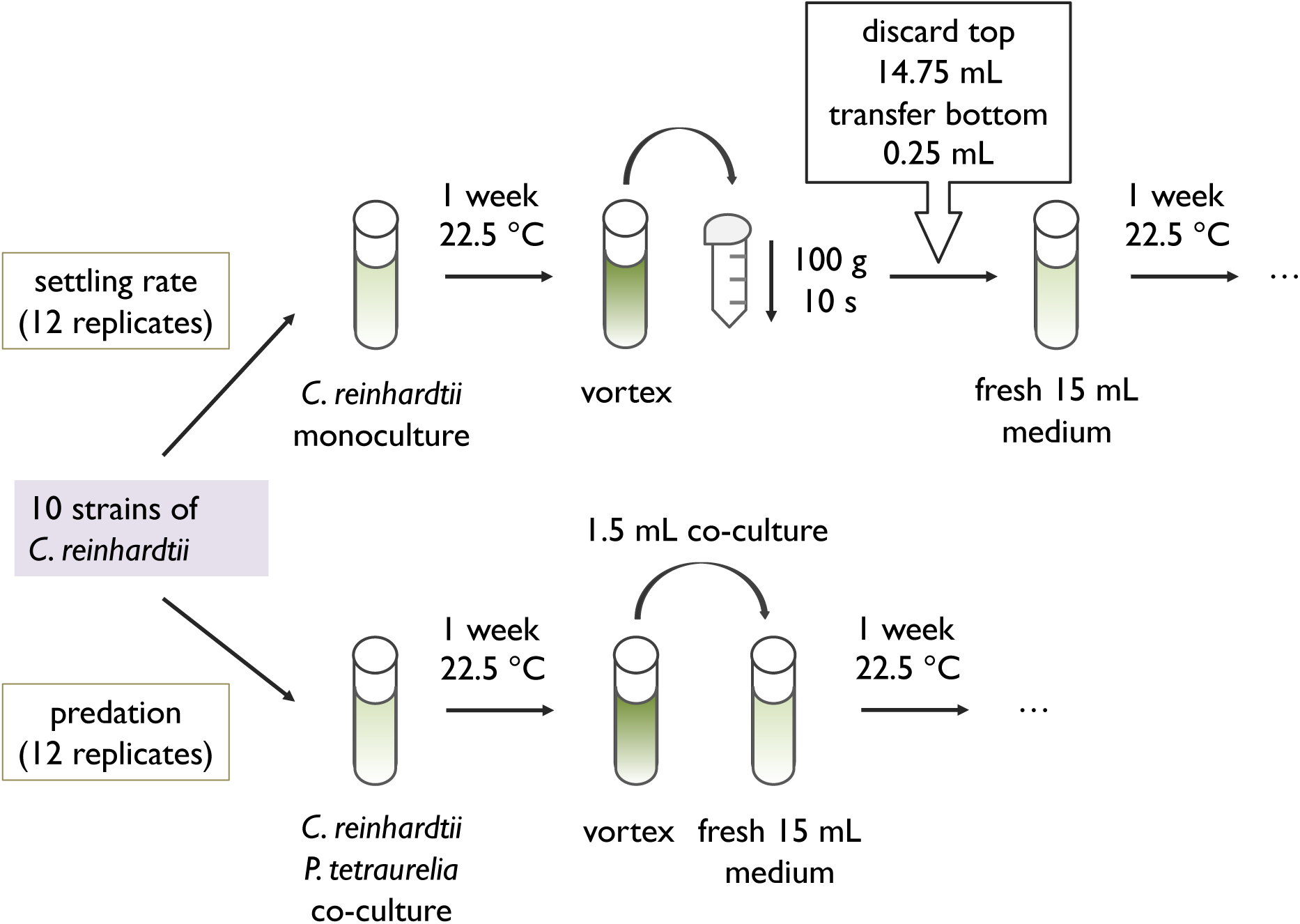
Experimental evolution diagram. A founder population consisting of 10 different strains of *C. reinhardtii*, all in equal proportion, was used to generate 24 replicate populations: 12 were subjected to settling selection that favors large cell clusters that settle rapidly; 12 were subjected to predation by the protist *P. tetraurelia*. Subsamples of each population were transferred to fresh medium every week, with approximately 15 generations elapsing between transfers, for a total of 40 transfers. For the settling rate selection, each population was transferred to a 15 mL Falcon tube and centrifuged at 100 x *g* for 10 s and only the bottom 0.25 mL was transferred to fresh medium. For populations under predation selection, 1.5 mL of co-cultured *C. reinhardtii* and *P. tetraurelia* were transferred to fresh medium every week.

Every two months (i.e., every eight transfers or 120 generations), 0.1 mL samples of each experimental population were diluted 100-fold and 30 μL was plated on 0.5% TAP agar to screen for multicellular isolates. Unicellular isolates are normally motile and form large colonies on 0.5% agar plates, whereas multicellular isolates are normally non-motile and form small colonies (preliminary data not shown). Three small colonies per population (presumably multicellular isolates labelled “A”, “B”, and “C”) were picked up using sterile toothpicks, inoculated into 2 mL of TAP medium in 24-well plates, incubated overnight and screened the next day for multicellular phenotypes by visual examination under light microscopy. Isolates that demonstrated multicellular phenotypes were diluted 10-fold, then 30 μL was replated on solid medium. The same procedure was repeated three times sequentially to ensure that each isolate was clonal and that multicellular phenotypes of isolates from the predation treatment were not transiently induced by the presence of the *P. tetraurelia* predator. All founder strains used to start the evolution experiment and evolved multicellular isolates were cryopreserved at −80°C using the GeneArt Cryopreservation Kit for Algae (Invitrogen, Thermal Fisher Scientific).

### Cells per cluster in the multicellular isolates

At the end of our evolution experiments we obtained pure culture isolates of several multicellular strains. To understand how cluster size distribution patterns varied among strains over a 6-day incubation period as well as the maximum number of cells per cluster, we subjected each strain to epifluorescence microscopy. Cells of evolved strains and their ancestors were scraped from TAP agar plates (3-4 inoculation loops, 0.09 cm in diameter), inoculated into 15 mL of TAP medium then cultured for five days to high density (approximately 6-8 x 10^6^ cells/mL). Next, we transferred cultures of each strain into 15 mL of TAP medium at a starting density of 10^4^ cells/mL, cultured on a 14:10-hour light:dark cycle at 22.5 °C, and sampled every 24 hours over the course of 6-days. We used TAP medium here because it is a standard medium commonly used in *Chlamydomonas* research (45); multicellular phenotypes were stable in TAP. Subsets of cell cultures were removed from well-mixed test tubes and transferred into 1.5 mL Eppendorf tubes (from day 1 to day 6, the amounts of cell cultures transferred were 1200, 900, 400, 300, 200, 100 μL). Samples were centrifuged at 14,000 × *g* for 1 minute and the supernatant was replaced by a 50% v/v ethanol-water solution to fix cells and render cell membranes permeable to the fluorescent nucleotide stain DAPI (4′,6-diamidino-2-phenylindole). After a 10-minute fixation period, samples were centrifuged again at 14000 × *g* for 1 minute and the supernatant replaced with distilled water containing 1 μg/mL DAPI. Cells stained with DAPI were kept in darkness for one hour at 25°C then moved to a 4°C refrigerator without light to prevent photobleaching until being imaged on an inverted epifluorescence microscope (the Eclipse Ti series, Nikon).

Prior to imaging, samples were removed from the refrigerator and gently pipetted. A 10 μL aliquot of each sample was mounted on glass microscope slides, then multiple images were recorded at 20x magnification using bright field and the DAPI filter set (Excitation/Emission: 358/461 nm) overlapping on top of each other. Images were taken of at least 20 clusters or cells per sample, providing a robust dataset for computing the distribution of cluster sizes within each population at a given time point. Next, images were manually screened to demarcate cluster boundaries and record cell number within each boundary using Cell Counter in ImageJ (https://imagej.nih.gov/ij/). Cell number was estimated by counting DAPI stained nuclei confined within cluster boundaries (e.g., **Figure 2**). This procedure was repeated four times for each of the evolved strains and twice for the ancestral strains.

**Figure 2.**
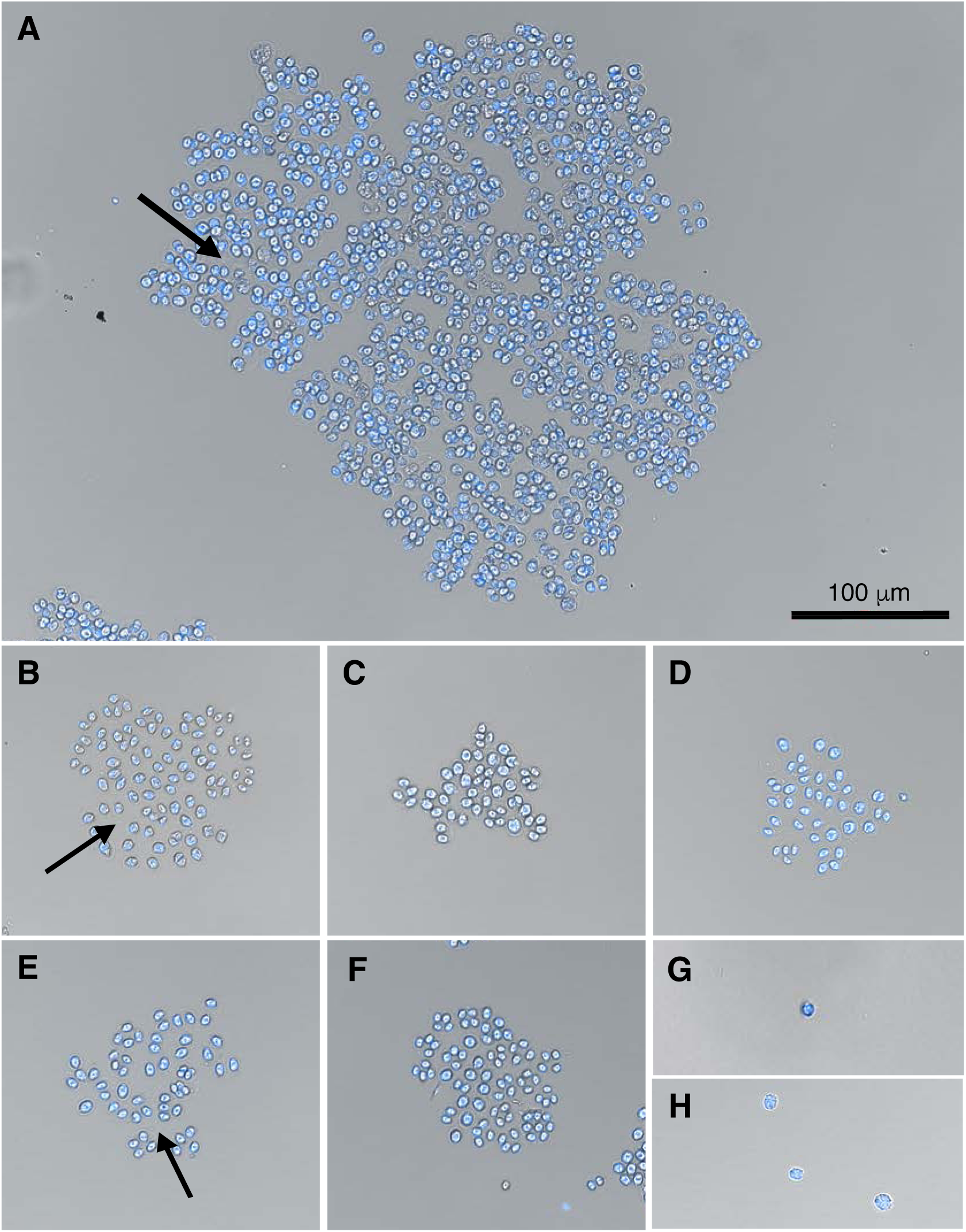
Microscopic images of each evolved strain and their unicellular ancestors stained by DAPI. (A) PS09-32-A; (B) PS11-32-B; (C) PS02-40-A; (D) PS02-40-B; (E) PS02-40-C; (F) PS12-40-C; (G) CC-2936; (H) CC-1690. Black arrows indicate examples of extracellular matrix. Genomic data indicate that strain PS12-40-C evolved from CC-1690 while the rest of multicellular strains evolved from CC-2936 (see **Table 3**).

### Modes of reproduction in multicellular isolates

Multicellularity can arise either by many cells aggregating (47, 48) or by one cell reproducing clonally and its daughter cells remaining intimately associated (49, 50). In order to distinguish between these two forms of multicellularity, ancestral and evolved strains were imaged using time-lapse transmission microscopy over 72 hours of growth. As before, cells were scraped from TAP agar plates (3-4 inoculation loops, 0.09 cm in diameter), inoculated into liquid TAP medium and cultured for five days to high density (approximately 6-8 x 10^6^ cells/mL).

Then, each culture was vortexed, transferred 1:100 into fresh TAP medium, and allowed to grow for two more days. Exponentially growing cultures were first mixed by vortexing, then diluted 1:10 in TAP medium, whereafter 100 μL from each culture was inoculated into the wells of a 96-well tissue culture plate containing100 μL of TAP medium per well. Well positions for each strain were determined by a random number generator. Wells not inoculated with algae were filled instead with 100 μL of TAP cell-free medium. For time-lapse microscopy, the 96-well plate was imaged at 200x magnification using the Nikon Eclipse Ti inverted microscope, where the field of view was positioned on cells near the center and at the bottom of wells. The time-lapse was run for 72 hours, capturing images of each well every 30 minutes. For each strain, five technical replicates were performed on the 96-well plate.

### Identifying the ancestors of evolved multicellular isolates

MATING TYPE ASSAY: To ascertain whether certain ancestral strains were more likely to evolve multicellularity than others, we identified alleles that differed among founder strains (such that each strain has a specific combination), and used those alleles as markers to determine the probable ancestor(s) of evolved multicellular strains. We first established the alleles at the mating type locus in evolved isolates, as the mating types of the founder strains had been previously identified. Each evolved isolate was subjected to PCR-amplification using either the *mid* or *fus1* locus (the *mid* gene is unique to mating type minus (mt-) cells, while the *fus1* is unique to mating type plus (mt+) cells). The *mid* locus was amplified using the primer pair MTM1F (CTG CTG GTA CAA AGG TGT GGC ACG) and MTM2R (CAT GCA GTC TCT CTC ACC CAT TCG GC), which were expected to produce a 750bp piece of the *mid* gene; the *fus1* locus was amplified using the primer pair MTP2F (GCT GGC ATT CCT GTA TCC TTG ACG C) and MTP2R (GCG GCG TAA CAT AAA GGA GGG TCG), which were expected to produce a 434bp piece of the *fus1* gene (51). After identifying the mating type of evolved isolates, we next compared how their whole genome sequences differed from the founder strains, giving us deeper insight into which ancestors produced multicellular isolates.

WHOLE GENOME SEQUENCING: Genomic DNA from evolved isolates and the ancestral strain CC-1690 (a wild-type strain in our lab) was extracted, using the *Chlamydomonas* DNA extraction protocol recommended by the Joint Genome Institute (https://www.pacb.com/wp-content/uploads/2015/09/Experimental-Protocol-DNA-extraction-of-Chlamydomonas-using-CTAB-JGI.pdf). Genomic DNA libraries were prepared, and samples were sequenced at 2 x 151bp length using the Illumina NextSeq platform at Microbial Genome Sequencing Center (MiGS, https://www.migscenter.com/). For the founder strains other than CC-1690, their sequenced reads were downloaded from NCBI. The quality of sequenced reads of evolved strains and founders was examined by FastQC and trimmed using Trimmomatic-0.36 (*options*: HEADCROP:16 LEADING:30 TRAILING:28 SLIDINGWINDOW:4:15 MINLEN:54). Paired reads were mapped to the Joint Genome Institute v5.5 assembly of the *C. reinhardtii* CC503 mt+ genome (52) using BWA-MEM default setting (53). Genomic variants (small polymorphisms such as SNPs (single nucleotide polymorphisms), indels (insertions and deletions), MNPs (multi-nucleotide polymorphisms) and complex events (composite insertion and substitution events)) were detected using FreeBayes v1.0.2-29 (*options*: -p 1 -m 3 -q 1) (54). Variants with quality scores greater than 20, read depth greater than 5X, and allele frequencies greater than 0.85, were analyzed. According to a previous study (44), each founder strain contains a genome fingerprint that distinguish them from each other (e.g., see Flowers et al. 2015 **Figure 2B**). Our approach was to generate a filtered genetic variant data set that consisted of nonreference variants observed in all strains together, and identify ancestral identities of multicellular isolates using strain-specific variants along with mating types in founder strains. Sequencing data were deposited at the NCBI Sequence Read Archive with project number PRJNA820479 and accession numbers SAMN27009621-4.

## Results

### Multicellular structures evolved under predation-selection but not under settling selection

A founder population of *C. reinhardtii* was created by mixing in equal proportion each the 10 strains described in **Table 1**; subsamples of this founder population were subjected to 12-fold replicated experimental evolution according to the scheme depicted in **Figure 1**. Hereafter, we will refer to <u>P</u>redation <u>S</u>election lines as PS01-PS12 and <u>S</u>ettling <u>R</u>ate <u>S</u>election Lines as SRS01-SRS12. Under the predator selective regime, after 16 weekly transfers we observed multicellular structures in two of the 12 replicate populations, PS05 and PS11. However, evolved isolates from those two populations did not maintain their multicellular phenotypes after serial plate transfers in the laboratory; neither did multicellular morphologies persist in these two populations in the following screen after eight weeks. After 32 weekly transfers, multicellular structures appeared in three populations, PS09, PS10 and PS11; one evolved strain was isolated from each population (PS09-32-A, PS10-32-B and PS11-32-B respectively). Other than PS10-32-B from PS10, the multicellular strains that evolved in populations PS09 and PS11 after 32 weekly transfers maintained multicellular morphologies after being plated on solid medium three times in succession, as well as in the experiments described below (**Figure 2: Panels A-B**). After an additional 8 weekly transfers (40 transfers altogether), multicellular structures were detected in two other predation-selected populations, PS02 and PS12; three evolved strains were isolated from PS02 (PS02-40-A, PS02-40-B, PS02-40-C) and one strain was isolated from PS12 (PS12-40-C). Each of the evolved strains isolated from PS02 and PS12 after 40 weekly transfers maintained their multicellular structures after being plated on solid medium three times in succession, as well as in the experiments described below (**Figure 2**: **Panels C-F)**. Together these observations indicate that the evolved multicellular phenotypes under predation selection are stably heritable. Under the settling rate selective regime, we observed multicellular structures in two of the 12 replicate populations, SRS05 and SRS 07, after 16 weekly transfers, and again in populations SRS03, SRS08 and SRS11 after 32 weekly transfers. However, isolates from those populations did not maintain their multicellular phenotypes after serial plate transfers in the laboratory. Thus, heritably stable multicellular structures were never observed in any of the settling rate selected populations over the course of 40 weekly transfers, or ∼600 generations.

In the unicellular ancestors, most cells exist as single cells in their respective populations (**Figure 2**: **Panels G-H**). By contrast, within each multicellular structure of the evolved strains, cell walls are visible surrounding individual two, four or eight cells, and these are held in place by transparent extracellular matrix (**Figure 2**: **Panels A-F**). Most evolved strains form clusters approximately 20-100 µm in size, except PS09-32-A, which can form clusters whose diameters range up to several hundred micrometers (**Figure 2**: **Panel A**). Development in the evolved multicellular strains is strictly clonal. Time-lapse microscopy over 72 hours shows that, compared to the ancestors where dividing cells only transiently form clusters before daughter cells are released (e.g., CC-2936, **Supplemental Video 1**), multicellular isolates develop by parental cells dividing into daughter cells within clusters over multiple rounds of cell division (**Supplemental Videos 2-7**). Because cells within each multicellular cluster are genetically identical, clusters likely function as units of selection (55, 56) under predation selection.

### Cells per cluster in the multicellular isolates

In order to determine cluster size distributions for evolved strains, we sampled, stained, and imaged growing cultures every 24 hours over a 6-day period, which approximated the serial transfer interval of our evolution experiments. Four biological replicates were performed for each of the evolved strains, and two biological replicates were performed for the ancestral strains. As distributions tended to be right-skewed, data were represented as boxplots that depict clusters’ central tendencies. Large outlier clusters were occasionally present in each population. These outliers, defined as data points that exceeded a distance of 1.5 times the interquartile range (IQR) above the 3^rd^ quartile, were excluded from further analyses.

We observed that multicellular cluster size varied among evolved strains over the course of 6-days. The median cluster size of multicellular strain PS09-32-A was >80 cells on day-1 and then maintained at >100 cells throughout the growth cycle, with a maximum of 156 cells on day-6 (**Figure 3A**). By contrast, median cluster sizes of PS11-32-B, PS02-40-A, PS02-40-B, and PS12-40-C ranged between 2 and 10 cells, with maximum sizes being reached on day-1 (6, 10, 7 and 7 cells, respectively) then gradually decreasing thereafter (**Figure 3B, C, D, F**). The median cluster size of strain PS02-40-C remained at 4 cells between day-1 and day-5 before declining to 2 cells on day-6 (**Figure 3E**). Median cluster sizes of the ancestral strains (CC-2936 and CC-1690, see the following paragraph for identification details) remained one cell per cluster over growth cycle, with multicellular clusters occurring occasionally during reproduction (**Figure 3G-H**).

**Figure 3.**
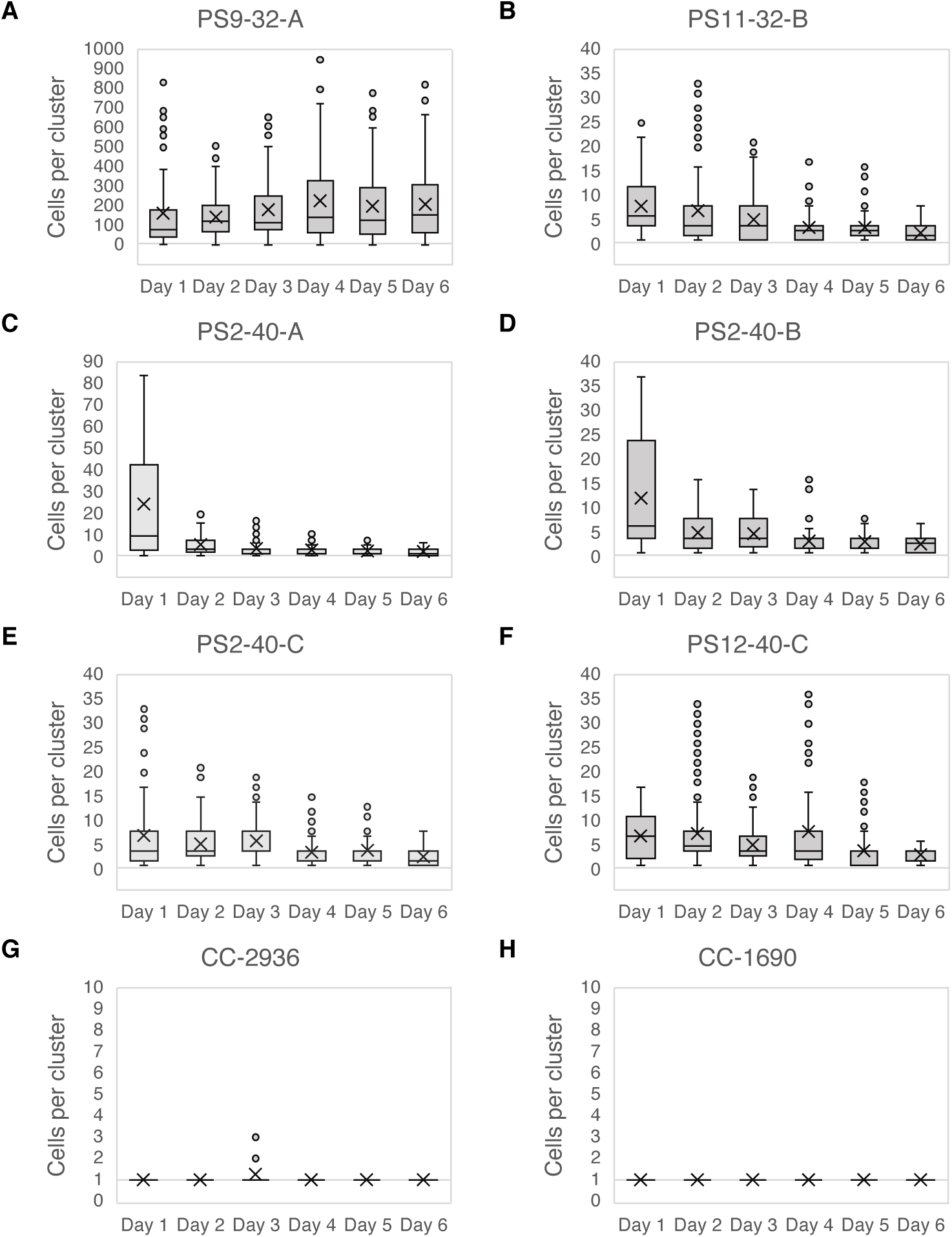
Box plots of cells per cluster over a 6-day period. Cluster sizes were measured by sampling cultures over six days of growth, staining nuclei with DAPI, and imaging using fluorescent microscopy. (A-F) Cluster sizes in the evolved strains; (G-H) Cluster sizes in the ancestors. An x or cross represents the mean of cells per cluster; the crossbar indicates the median of cells per cluster. The grey horizontal bar in each figure represents one cell per cluster. Note the y-axis scales vary among strains.

We found that there was a significant difference among evolved strains and ancestral strains in the median cluster sizes (Kruskal-Wallis Test, H = 242.22, df = 7, p << 0.01). To ensure our comparisons were of similar developmental stages, the median for each strain was measured at the time point at which it reached its maximum value, as in Herron, Borin (42). Particularly, the median cluster size of PS09-32-A (156 cells) was significantly larger than those of other multicellular strains (6, 10, 7, 4 and 7 cells from PS11-32-B, PS02-40-A, PS02-40-B, PS02-40-C, PS12-40-C, respectively) and those of their ancestors (1 and 1 cell from CC-2936 and CC-1690) (**Table 2**). There were no significant differences in median cluster sizes among all the multicellular strains other than PS09-32-A (**Table 2**).

**Table 2.**
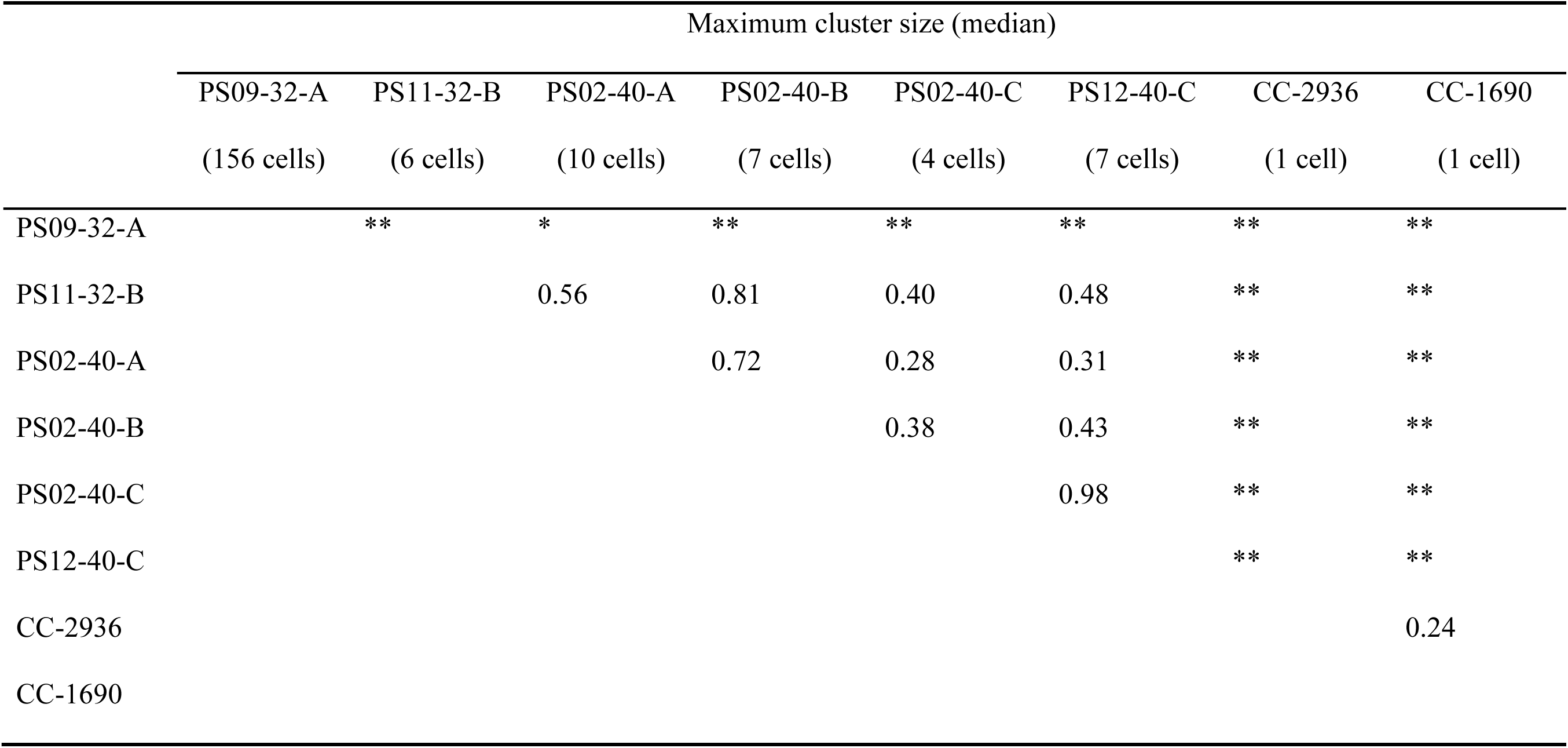
P-values for Dunn’s Test for all pairwise comparisons between the maximum cluster sizes in different strains. Single asterisks indicate p-values of <0.05. Double asterisks indicate p-values of < 0.01.

### Multicellular isolates evolved from the founder strains CC-2936 and CC-1690

PCR analysis of mating types in the evolved multicellular isolates revealed that all were mt+, suggesting they originated from at least one of the mt+ founder strains (**Supplemental Figure 1**). Therefore, for the following genomic analysis, we only compared evolved isolates with mt+ founders to identify which ancestor genotype(s) they were most likely to have originated from.

We performed whole genome sequencing of PS09-32-A, PS11-32-B, PS02-40-A in PS02 and PS12-40-C to an aligned sequencing depth of ∼15X, 15X, 14X and 18X using paired end (2 x 151bp) Illumina sequencing. One of the mt+ founder strains, CC-1690, was re-sequenced for a separate study (not published) to an aligned depth of 44X using the same Illumina sequencing method. We did not sequence PS2-40-B and PS2-40-C in PS02 owing to the fact that they are phenotypically similar to PS02-40-A, and are likely the same genotype as PS2-40-A, or only differ by a few mutations, as they were isolated from the same population. For the remaining four mt+ founder strains, their sequenced reads were downloaded from NCBI using accession numbers SAMN03272879, SAMN03272875, SAMN03272877 and SAMN03272872 for strains CC-2937, CC-2936, CC-2344 and CC2343, respectively (44). Based on the sequence alignments to ∼112 Mb of the *C. reinhardtii* CC-503 mt+ reference genome (52), we generated a filtered genetic variant data set that consisted of 4,714,560 nonreference variants in all strains together. Each strain contains a genome fingerprint based on the location of their genetic variants across 17 *C. reinhardtii* chromosomes. By visually comparing the genome-wide distributions of those variants and through direct comparisons, we can infer the most likely ancestors of evolved multicellular isolates in the populations PS02, PS09, PS11 and PS12.

Among mt+ founder strains, lab strain CC-1690 is diverged from reference strain CC-503 by 90,242 variants in our filtered data set, while the field isolates CC-2937, CC-2936, CC-2344 and CC-2343 differ from CC-503, respectively, by 1,651,964, 1,194,250, 2,102,758, 2,187,600 variants in 111,098,438 sites across the reference genome (**Table 3**). Among evolved strains, PS09-32-A, PS11-32-B and PS02-40-A contain 1,169,953, 1,110,266 and 1,144,326 nonreference variants, respectively, while PS12-40-C only contains 77,708 variants (**Table 3**). When comparing the genome-wide distributions of nonreference variants between mt+ founders and evolved strains, we found that the distributions in PS09-32-A, PS11-32-B and PS02-40-A visually match to that in CC-2936 while the distribution in PS12-40-C visually match to that in CC-1690 (**Figure 4**). When directly comparing the variants between mt+ founders and evolved strains, we found that > 99% of variants identified in the evolved strains PS09-32-A, PS11-32-B and PS02-40-A matched to the founder strain CC-2936, while 100% of the variants identified in PS12-40-C matched to the founder strain CC-1690 (**Table 3**), suggesting that PS09-32-A, PS11-32-B, PS02-40-A evolved from CC-2936 and that PS12-40-C evolved from CC-1690. The < 1% of variants in PS09-32-A, PS11-32-B, PS02-40-A that did not match to CC-2936 is likely due either to different sequencing methods (e.g., CC-2936 was previously sequenced at 2 x 51bp length using the Illumina HiSeq 2000 sequencer (44) while the evolved strains here were sequenced at 2 x 151bp length using the Illumina NextSeq 2000 sequencer), or to mutations that accumulated in these lines since they were sequenced by Flowers, Hazzouri (44).

**Table 3.** Comparisons between genomic variants in evolved and mt+ founder strains. The numbers in parenthesis after strain names indicate the number of nonreference variants identified in each strain (see Methods). Heat maps are color-coded (light grey/low percentage match < grey/middle percentage match < dark grey/high percentage match).

|  | CC-1690 | CC-2937 | CC-2936 | CC-2344 | CC-2343 |
| --- | --- | --- | --- | --- | --- |
|  | (90,242) | (1,651,964) | (1,194,250) | (2,102,758) | (2,187,600) |
| PS09-32-A<br>(1,169,953) | 59.56% | 35.46% | 99.93% | 22.50% | 18.41% |
| PS11-32-B<br>(1,110,266) | 59.15% | 35.11% | 99.90% | 22.28% | 18.18% |
| PS02-40-A<br>(1,144,326) | 59.59% | 35.40% | 99.93% | 22.42% | 18.32% |
| PS12-40-C<br>(77,708) | 100% | 2.02% | 3.58% | 1.33% | 1.09% |

**Figure 4.**
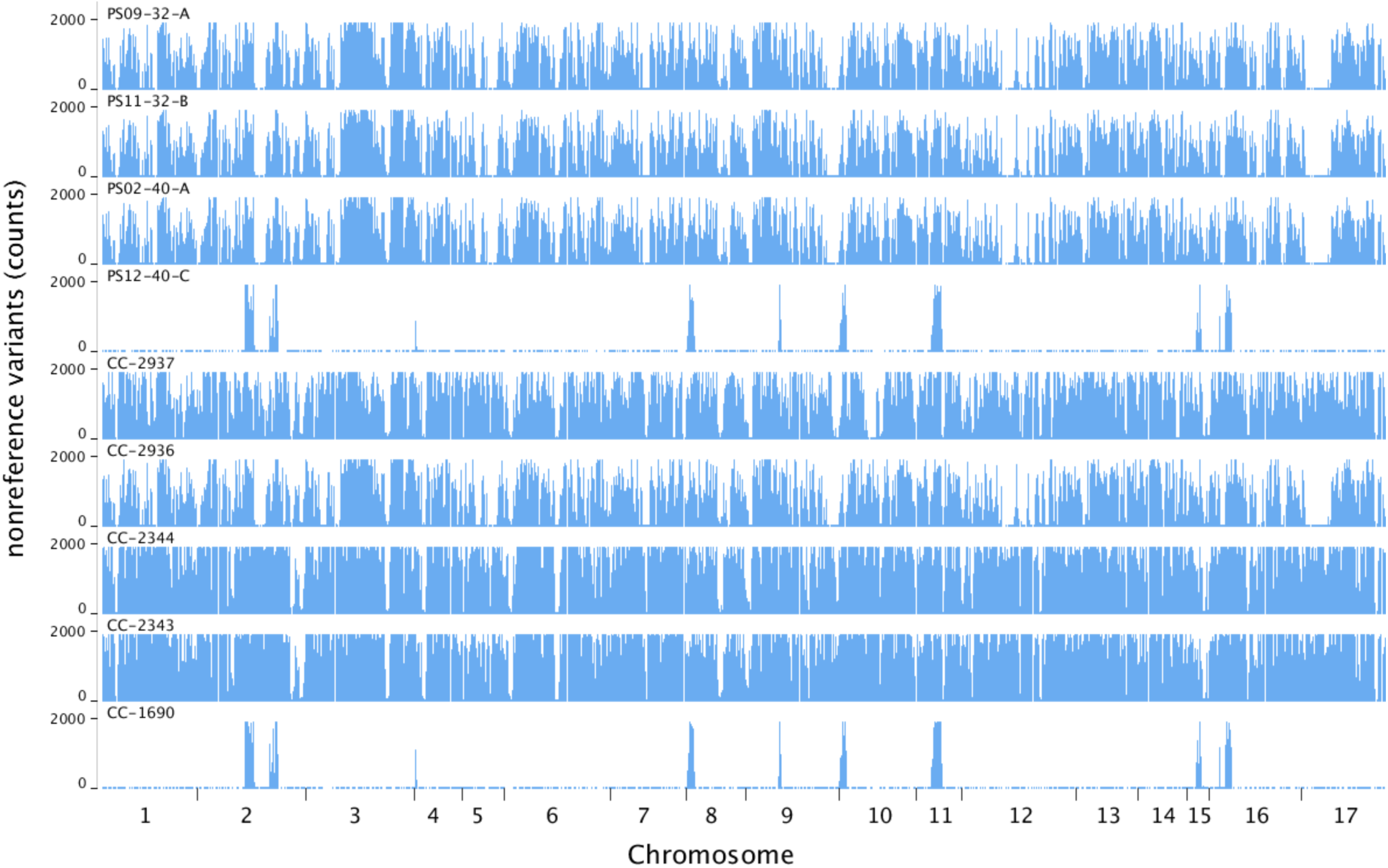
Genome-wide distributions of nonreference variants in evolved and mt+ founder strains on 17 *Chlamydomonas* chromosomes.

To further illustrate that specific allele combinations in the founder strains CC-2936 and CC-1690 are also present in their respective evolved isolates, **Table 4** lists 25 loci across different chromosomal locations that differ among five mt+ founders. The nucleotides in those 25 positions in PS09-32-A, PS11-32-B and PS02-40-A all match to the ones in CC-2936, while the nucleotides in those positions in PS12-40-C all match to CC-1690 (**Table 4**), indicating that CC-2936 is the ancestor of PS09-32-A, PS11-32-B and PS02-40-A and that CC-1690 is the ancestor of PS12-40-C.

**Table 4.** Examples of genomic variants that differ among mt+ founder strains and their presence and absence in the evolved strains.

| Strain | CC-1690 | CC-2937 | CC-2936 | CC-2344 | CC-2343 | PS09-32-A | PS11-32-B | PS02-40-A | PS12-40-C |
| --- | --- | --- | --- | --- | --- | --- | --- | --- | --- |
| Position |  |  |  |  |  |  |  |  |  |
| Chrom_1 1782815 | A | T | T | T | T | T | T | T | A |
| Chrom_1 1796517 | A | G | A | A | A | A | A | A | A |
| Chrom_2 531304 | G | G | A | G | G | A | A | A | G |
| Chrom_2 906332 | T | T | T | C | T | T | T | T | T |
| Chrom_2 1216720 | C | C | C | C | T | C | C | C | C |
| Chrom_3 666196 | C | A | T | A | A | T | T | T | C |
| Chrom_3 983155 | AA | AG | GG | GG | AG | GG | GG | GG | AA |
| Chrom_3 1128242 | C | C | T | C | C | T | T | T | C |
| Chrom_4 14191 | TGA | TGA | TGA | TA | TGA | TGA | TGA | TGA | TGA |
| Chrom_4 236504 | G | G | G | G | A | G | G | G | G |
| Chrom_5 763807 | G | A | A | A | A | A | A | A | G |
| Chrom_5 888052 | C | T | C | C | C | C | C | C | C |
| Chrom_5 1624406 | G | G | A | G | G | A | A | A | G |
| Chrom_6 122423 | ACG | ACA | ACA | ATG | ACG | ACA | ACA | ACA | ACG |
| Chrom_6 204099 | T | T | T | T | C | T | T | T | T |
| Chrom_7 1392564 | CCCGC | CC | CC | CC | CC | CC | CC | CC | CCCGC |
| Chrom_8 301955 | TG | AA | TG | TG | TG | TG | TG | TG | TG |
| Chrom_8 533535 | A | A | G | A | A | G | G | G | A |
| Chrom_9 133015 | A | A | A | C | A | A | A | A | A |
| Chrom_10 69703 | C | C | C | C | T | C | C | C | C |
| Chrom_11 448221 | A | C | C | C | C | C | C | C | A |
| Chrom_13 679947 | ATCTAT | AAC | ATCTAT | AAC | AAC | ATCTAT | ATCTAT | ATCTAT | ATCTAT |
| Chrom_14 153285 | G | G | A | G | G | A | A | A | G |
| Chrom_15 135507 | C | C | C | T | C | C | C | C | C |
| Chrom_16 312391 | C | C | C | C | G | C | C | C | C |

## Discussion

The emergence of multicellular organisms and the emergence of eukaryotes are widely regarded as major evolutionary transitions in the history of life (57). However, unlike eukaryogenesis, which seems to have been a singular event (58, 59), multicellularity has arisen at least 25 times across widely divergent lineages (20). Within the volvocine algae, multicellularity arose in some lineages as recently as the Jurassic and Cretaceous periods (126-49 million years ago) (37, 60–62). Still, even though multicellularity has arisen repeatedly and in diverse lineages, little is known about the relative roles that stochastic versus deterministic factors play in driving this transition. Previous work has shown that in the lab simple, obligate multicellularity can evolve in the unicellular green alga *C. reinhardtii* under either predation or settling selection (41, 42). Here, we used experimental evolution to assess the influences that genetic ancestry and selective environment have on whether single cells become stably multicellular. We established a founder population consisting of 10 different strains of *C. reinhardtii*, all in equal proportion.

This founder population was used to generate 24 replicate populations: 12 were subjected to predation by the protist *P. tetraurelia*, 12 were subjected to settling selection that favors large cell clusters that settle rapidly. After ∼600 generations, heritable, obligate multicellular structures arose in four of 12 predation-selected populations, but never in the settling rate selected populations. Furthermore, we found that multicellular isolates evolved from only two founders CC-1690 and CC-2936, and that the size and structure of some multicellular isolates from the founder CC-2936 differed from each other. Whether the obligate multicellularity we observed was at a stable point or at some point along a continuum of evolving multicellularity, our results indicate that both deterministic and stochastic factors play significant roles on the trajectories by which populations evolve multicellularity.

If the evolutionary transition to multicellularity was purely deterministic, we would expect that similar selective environments would result in similar phenotypes regardless of ancestral genotypes. However, the fact that we observed different types of multicellularity evolve under predation selection, even from the same ancestor (CC-2936), suggests that stochastic processes like the accumulation of random mutations can lead to largely different outcomes. That being said, the large clusters formed by PS09-32-A (> 100 cells most of time) can be costly, as the growth of a colony may be limited because interior cells may not have the same access to nutrients or environmental signals (e.g. oxygen or lights) compared to exterior cells (63, 64). If this is true, then it is possible that PS09-32-A might have evolved a smaller cluster size similar to what we observed in other multicellular isolates had we allowed the experiment to run longer. Such a response to selection would demonstrate that evolutionary process is still deterministic at the phenotypic level.

We also found that the multicellular isolates evolved in three of 12 predation-selected populations came from founder CC-2936, while the multicellular isolate evolved in the other predation-selected population descended from founder CC-1690. We did not observe multicellular isolates evolved from any other founders. These results suggest that particular suites of genetic variants had accumulated in those strains prior to our evolution experiments, and that these facilitated the transition to multicellularity in our selective regimen. An inverse question to ask is: why are some founders more resistant than others to selection pressures expected to favor evolution of multicellularity? Were the populations quickly purged of certain founders just because they grew less quickly or to a lower stationary phase density? In other words, were the results biased at the onset because certain strains have poor competitive ability, relative to others? To test these ideas, future work should be directed toward competition experiments among founders, and/or similar evolutions using the same founders where populations are sequenced at different time points to monitor changes in the frequency of allelic markers by which founders can be differentiated.

Multiple reasons could explain why multicellularity did not evolve under our settling selection regimen. First, in our experiments settling rate selection was episodic, occurring only once a week, whereas predation selection was continuous. Second, our settling rate selection (centrifugation at 100 *g* for 10 s, with pelleted cells transferred to fresh medium once a week) differed slightly from that of Ratcliff, Herron (41) (100 *g* for 5 s, with pelleted cells transferred every 3 days). Third, so that the two treatments could be compared it was essential that the settling selection and the predation experiments be carried out in same medium. We elected to use COMBO, a defined freshwater medium for co-culturing algae and zooplankton (46) that had been used successfully in our earlier predation experiment (42). Significantly, COMBO does not contain an organic carbon source, therefore, it only supports photoautotrophic growth of *C. reinhardtii* (65). In this key respect, COMBO differs from the TAP medium more commonly used in *Chlamydomonas* research (including the Ratcliff, Herron (41) settling selection experiment). TAP contains acetate, which supports heterotrophic growth by *C. reinhardtii*.

COMBO necessarily brings about a different physiological state than TAP, and this difference may influence *C. reinhardtii*’s latent tendency to form clonal assemblages and/or extracellular matrices that cause bodies to settle rapidly in aqueous media. Fourth, by luck of the draw we might not have initiated our experiment with the right set of ancestral genotypes, or with sufficient standing genetic variation. Ratcliff, Herron (41) initiated their evolutions using an outbred population that had a high level of standing genetic variation; moreover, all populations went through one round of sexual reproduction (within-population mating) during the evolutions. After 73 transfers (∼315 generations) with settling rate selection imposed every three days, multicellular *C. reinhardtii* evolved in only one of 10 replicate populations. The difference between our results and those of Ratcliff, Herron (41) suggests that the amount of standing genetic variation might play a role in the tempo and mode by which multicellularity emerges.

Finally, perhaps multicellularity did not evolve under the settling selection here simply because the outcome is stochastic. After all, in Ratcliff, Herron (41) only one of 10 replicate populations evolved multicellularity under settling selection. If we use this as an estimate of the probability of multicellularity evolving under settling selection, the binomial probability of zero of twelve replicates evolving multicellularity would be 0.28 (p(x=0) given 12 trials, each with a probability of success of 0.1), far too likely to rule out.

Regarding the genetic basis of multicellularity in our experiments, we only sequenced evolved isolates exhibiting multicellularity at low coverages (14-18X), as our objective was to determine ancestry. Evolved unicellular isolates, in particular ones sharing a common ancestor with the evolved multicellular isolates, were not sequenced due to budget constraints. Therefore, comparisons between unicellular and multicellular evolved isolates, or between founder strains and their evolved multicellular isolates, are not possible for the investigation on the mutational trajectories to multicellularity here. Future directions include repeated backcrossing multicellular traits into the founder or wild-type strains to generate near isogenic lines, or directly sequencing the candidate genes known to cause multicellular phenotypes to identify the mutation(s) responsible for the transitions to multicellularity. For example, previous work has shown that the retinoblastoma (*RB*) cell cycle regulatory pathway is involved in cluster formation in the alga *Gonium*, an undifferentiated colonial relative of *Chlamydomonas* (33). Indeed, expression of the *Gonium RB* gene in a *Chlamydomonas* strain lacking its *RB* gene caused this mutant to form colonial assemblies containing 2-16 cells (33). It is plausible that mutations involved in the cell cycle pathway such as *RB* may underlie the transition from unicellular to multicellular phenotypes in our evolution experiments.

Taken together, our findings show that past genetic changes in some strains of *Chlamydomonas* can shift the probabilities of alternative evolutionary paths towards multicellularity. Further, subsequent mutations can play a significant role in shaping divergent multicellular forms even in populations evolved under the same condition.

## Acknowledgements

This work was supported by NASA Astrobiology Institute Award NNA17BB05A (Rosenzweig PI, Herron co-I), NASA Exobiology 80NSSC20K0621 (Rosenzweig PI), and NASA ICAR 80NSSC23K1357 (Rosenzweig PI). The authors thank Ozan Bozdag and Alireza Zamani-Dahaj in the Ratcliff lab for useful discussion and for their assistance with epifluorescence microscopy. We also thank Peter Conlin, Ross Lindsey, and Emily Cook for their helpful comments on earlier drafts of this manuscript. This material is based upon work while MDH was serving at the National Science Foundation.

## Notes

### Competing Interest Statement

The authors have declared no competing interest.

### Summary of Updates

The acknowledgements of the manuscript has been revised to reflect the support from NASA Exobiology 80NSSC20K0621 (Rosenzweig PI) and NASA ICAR 80NSSC23K1357 (Rosenzweig PI), and to acknowledge the helpful comments from Peter Conlin, Ross Lindsey, and Emily Cook on earlier drafts of the manuscript.

