## Supplemental Figure 1 for "Genetic predisposition towards multicellularity in *Chlamydomonas reinhardtii*"

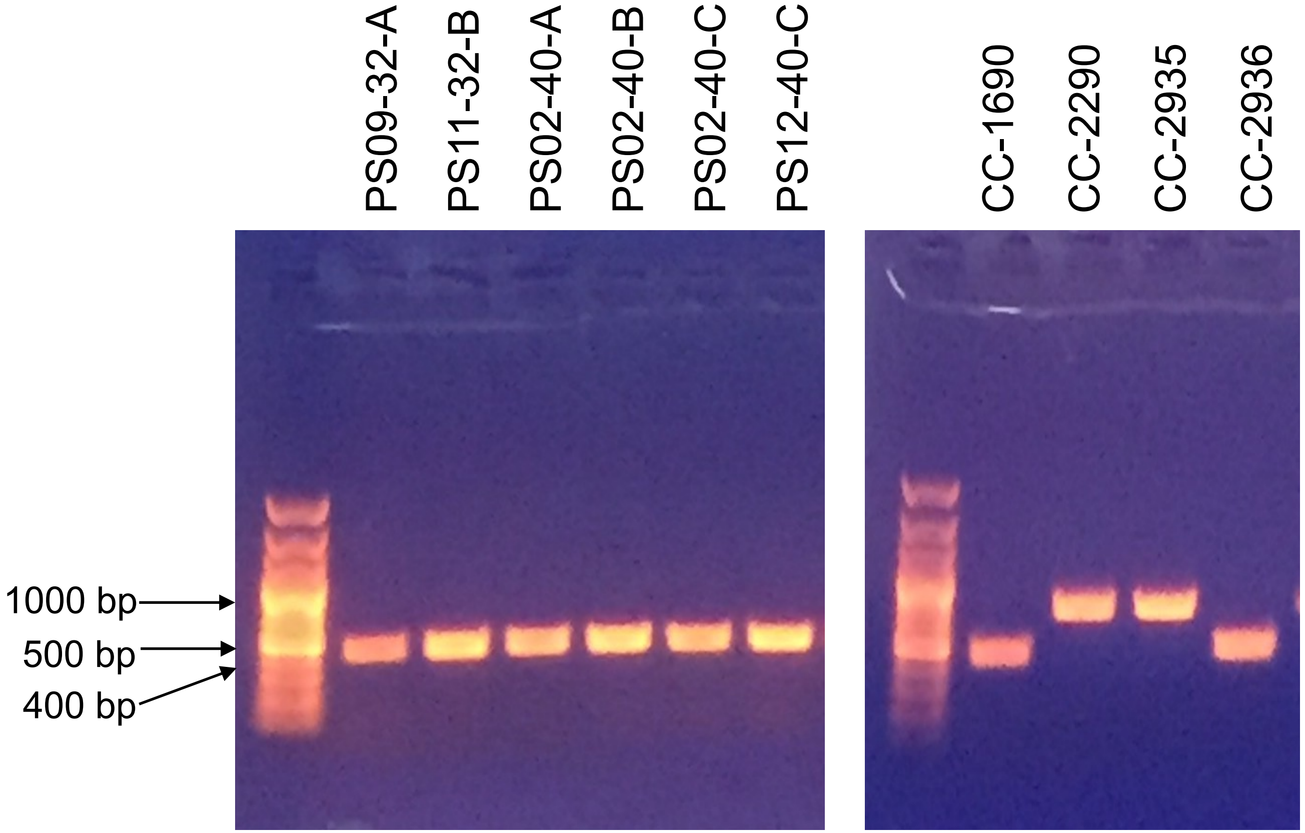


**Supplemental Figure 1.** Mating type PCR analysis of the evolved multicellular strains (PS09-32-A, PS11-32-B, PS02-40-A, PS02-40-B, PS02-40-C, PS12-40-C) and ancestral strains (CC-1690, CC-2290, CC-2935, CC-2936) with *mid*-specific primers (MTM1F and MTM2R) and *fus1*-specific primers (MTP2F and MTP2R).
